# Evaluation of pneumococcal serotyping in nasopharyngeal carriage isolates by latex agglutination, whole genome sequencing (PneumoCaT) and DNA microarray in a high pneumococcal carriage prevalence population in Malawi

**DOI:** 10.1101/2020.08.17.255224

**Authors:** Todd D. Swarthout, Andrea Gori, Naor Bar-Zeev, Arox W. Kamng’ona, Thandie S. Mwalukomo, Farouck Bonomali, Roseline Nyirenda, Comfort Brown, Jacquline Msefula, Dean Everett, Charles Mwansambo, Katherine Gould, Jason Hinds, Robert S. Heyderman, Neil French

## Abstract

**Background:** Accurate assessment of the serotype distribution associated with pneumococcal colonization and disease is essential for the evaluation and formulation of pneumococcal vaccines and informing vaccine policy.

**Methods:** We evaluated pneumococcal serotyping concordance between latex agglutination, PneumoCaT by whole genome sequencing (WGS) and DNA microarray using samples from community carriage surveillance in Blantyre, Malawi. Nasopharyngeal swabs were collected, following WHO recommendations, between 2015 and 2017, using stratified random sampling among study populations. Participants included healthy children 3–6 years old (PCV13 vaccinated as part of EPI), healthy children 5–10 years (age-ineligible for PCV13), and HIV-infected adults (18–40yrs) on ART. For phenotypic serotyping we used a 13-valent latex kit (SSI, Denmark). For genomic serotyping we applied PneumoCaT pipeline to whole genome sequence libraries. For molecular serotyping by microarray we used the BUGS Bioscience DNA microarray.

**Results:** 1347 samples were analysed. Concordance was 90.7% (95% CI: 89.0–92.2) between latex and PneumoCaT; 95.2% (93.9–96.3) between latex and microarray; and 96.6% (95.5–97.5) between microarray and PneumoCaT. By detecting carried vaccine serotype (VT) pneumococcus in low relative abundance (median 8%), microarray increased VT detection by 31.5% compared to latex serotyping.

**Conclusion:** All three serotyping methods were highly concordant in identifying dominant serotypes. Latex serotyping is accurate in identifying vaccine-serotypes and requires the least expertise and resources for field-implementation and analysis. However, WGS, which adds population structure, and microarray, which adds multiple-serotype carriage, should be considered at regional reference laboratories while investigating the importance of VT in low relative abundance in transmission and disease.

## Introduction

*Streptococcus pneumoniae* colonises the nasopharynx of healthy individuals. Although carriage is usually asymptomatic, nasopharyngeal (NP) colonization is a prerequisite for disease including otitis media, sinusitis, pneumonia, bacteraemia, and meningitis. (1) The pneumococcus is estimated to be responsible for over 318 000 (uncertainty ratio [UR]: 207 000–395 000) deaths every year in children aged 1 to 59 months, with the highest mortality burden among African children.(2) Evidence also shows that HIV-infected children and adults are at significantly higher risk of invasive pneumococcal disease (IPD) than their HIV-uninfected counterparts. (3, 4)

Current multivalent pneumococcal conjugate vaccines (PCV) target subsets of the nearly 100 capsular serotypes known to be expressed by the pneumococcus. PCV reduces nasopharyngeal carriage of the pneumococcal serotypes they contain, known as vaccine serotypes (VT). With reduced carriage among the vaccinated there is then reduced risk of VT-IPD (direct protection) and reduced transmission, therefore reduced risk of VT-IPD among those PCV-unvaccinated (indirect protection). However, non-vaccine serotypes (NVT) have the potential to fill the ecological niche, becoming more common in carriage and disease. (5–7) This phenomenon, known as serotype replacement, may be more pronounced in low-income settings because of higher prevalence, density and diversity of pneumococcal carriage, and represents a considerable risk to the global pneumococcal immunisation strategy.(8) Serotype distribution differs between continents as well as individual countries. (9) Given these differences, accurate assessment of the serotype distribution associated with both pneumococcal colonization and pneumococcal disease is needed in the evaluation, formulation and delivery of pneumococcal vaccines.

A pneumococcal serotyping method suitable for use in robust carriage and surveillance studies should therefore, at minimum, be accurate in its serotype assignment, particularly in relation to VTs. Additional desirable parameters include detection of most or all serotypes, ability to detect multiple serotypes in carriage (common in high burden settings (10, 11), more in-depth information on genotype, suitable to scale up for large projects, and practical for resource-poor settings. Unfortunately, work in resource-poor settings can too often limit the number of these parameters that can be achieved.

The gold-standard serotyping method, the Quellung reaction, was developed in the early 1900s and is performed by testing colonies with a set of type-specific antisera. (12) Bacteria are observed by microscopy, with serotype defined by observing apparent capsular swelling in reaction to the type-specific antisera. It is laborious, requires frequent use to maintain skills, requires a complete set of type-specific antisera, and is therefore mainly performed by reference laboratories. The PneuCarriage project, a large, multi-centre study, was established with the aim of identifying the best pneumococcal serotyping methods for carriage studies. (13) The Project identified microarray with a culture amplification step as the top-performing method. While robust and systematic, their decision algorithm did not take into account parameters such as cost, skill level and resources needed for assay implementation and maintenance, as well as output processing and interpretation.

Here we describe, in the context of an ongoing field-based study, (14) the level of concordance between three methods commonly used during ongoing routine pneumococcal surveillance activities in our work: latex agglutination, microarray and serotyping-by-sequencing (using the PneumoCaT pipeline). We also address parameters that researchers and policymakers can consider when deciding which assay to implement in their local setting.

## Materials and Methods

### Study Setting

Blantyre is located in Southern Malawi with an urban population of approximately 1.3 million.

### Study Population and Recruitment

Samples were collected as part of a larger 3.5-year pneumococcal carriage surveillance project, as described elsewhere. (14) In brief, this was a prospective rolling cross-sectional observational study using stratified random sampling to measure pneumococcal nasopharyngeal carriage in Blantyre, Malawi. Samples used in this analysis were collected during the first two years of twice-annual cross-sectional surveys, from June 2015 to April 2017. Recruitment included three groups: i) healthy children 3–6 years old who received PCV13 as part of routine EPI, ii) healthy children 5–10 years old who were age-ineligible to receive PCV13 as part of EPI, and iii) HIV-infected adults (18–40yrs) on ART.

### Sample Selection

For concordance analyses between the three methods, all samples were included that had serotyping results available from each of the three methods (latex, microarray, serotyping-by-sequencing). From the total nasopharyngeal swab (NPS) samples collected during the larger surveillance project (including 1,044 from children 3–6 years old [PCV-vaccinated], 531 children 5–10 years old [PCV-unvaccinated, age-ineligible] and 428 HIV-infected adults on ART), 1347 samples were culture-confirmed for *S. pneumoniae* and also had results available from the microarray and serotyping-by-sequencing. The final concordance analysis included 846 children 3–6 years old (PCV13-vaccinated), 422 children 5–10 years old (age-ineligible for PCV13 vaccination) and 79 adults (HIV-infected and PCV13-unvaccinated). (Figure 1) Sample selection for microarray and serotyping-by-sequencing was done independently and blind to latex serotype data.

**Figure 1.**
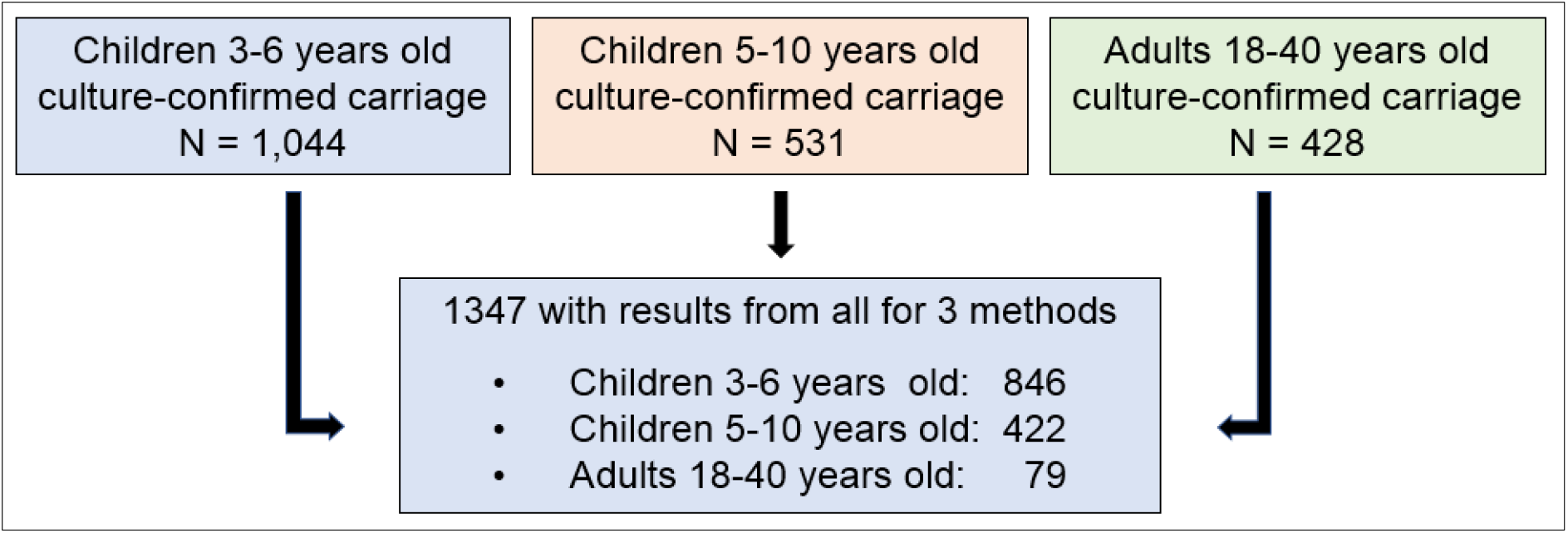
Sample selection for analysis. Sample were collected during 4 rolling cross-sectional surveys from June 2015 to April 2017. From the total nasopharyngeal swab samples collected 1347 samples had results available from the three assays under review. The method of selection for microarray and PneumoCaT was done independent of available latex serotype data.

### Nasopharyngeal Swab Collection

Collection of NP swabs is described elsewhere. (14) In brief, an NP swab sample was collected from each participant using a nylon flocked swab (FLOQSwabs™, Copan Diagnostics, Murrieta, CA, USA) and then immediately placed into 1.5mL skim milk-tryptone-glucose-glycerol (STGG) medium and processed at the Malawi–Liverpool–Wellcome Trust (MLW) laboratory in Blantyre, a ccording to WHO recommendations. (15) Samples were frozen on the same day at −80°C. (Figure 2)

**Figure 2.**
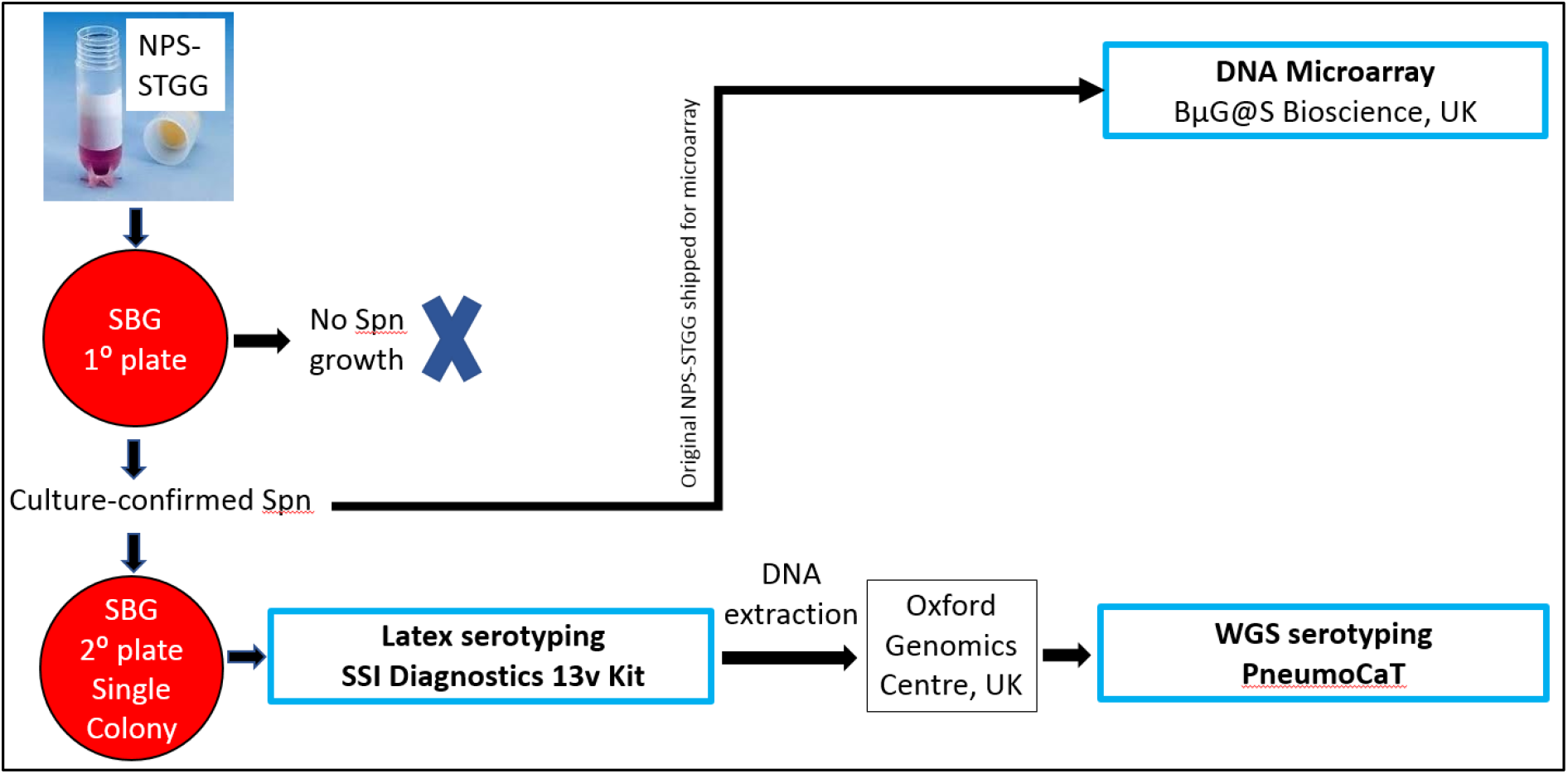
Laboratory procedures. Nasopharyngeal swabs (NPS) were inoculated into STGG and subsequently plated on a growth agar of sheep blood and gentamycin. Bacteria growth (from single-colony picks) from samples culture-confirmed for *Streptococcus pneumoniae* were used for latex serotyping. Remaining pure-growth isolates, retained at −80°C in sterile STGG, were later grown for DNA extraction and WGS. Aliquots of original samples (NPS-STGG) that were culture-confirmed for Streptococcus pneumoniae were assessed by microarray. NPS=nasopharyngeal swabs, STGG=skim-milk-tryptone-glucose-glycerol, WGS=whole-genome sequencing, Spn=*Streptococcus pneumoniae*, SBG=sheep blood and gentamycin, SSI= Statens Serum Institute, 13v=13-valent, NSP-STGG=NPS inoculated into STGG.

### NPS Culture for Pneumococcal Screening & Serotyping

30uL NPS–STGG was plated on a clean sheep blood-gentamicin (SBG; 7% SBA, 5 μl gentamicin/mL) agar plate (primary plate) and incubated overnight at 37°C in ~5% CO_2_. Plates showing no *S. pneumoniae* growth were incubated overnight a second time before being reported as negative. *S. pneumoniae* was identified by colony morphology and optochin disc (Oxoid, Basingstoke, UK) susceptibility. The bile solubility test was used on isolates with no or intermediate (zone diameter <14mm) optochin susceptibility. A single colony of confirmed pneumococcus was selected and grown on a clean SBG plate (secondary plate), following the same process as the primary plate. (Figure 1)

### Latex Serotyping

Pneumococcal growth from secondary plates was used for serotyping by latex agglutination (ImmuLex™ 7-10-13-valent Pneumotest; Statens Serum Institute, Denmark), following manufacturer guidelines. Using a reaction card and a sterile inoculation loop, a small sweep of an overnight bacterial culture was mixed with saline and a series of individual Pneumotest-Latex reagents in suspension. The card was rocked manually and observed for agglutination. A Pneumotest-Latex chessboard was used to determine which serotype is associated with the observed set of agglutination reactions. The kit allows for differential identification of each PCV13 VT (1, 3, 4, 5, 6A, 6B, 7F, 9V, 14, 18C, 19A, 19F, 23F). Other than for a limited number of serogroups (6, 7, 9, 18, 19, 23) for which the kit provides serogroup differentiation, there is no further differential identification of NVT serogroups or serotypes. NVT and non-typeable isolates were reported as NVT. Samples were batch-tested on a weekly basis, with technicians blinded to the sample source. After serotyping was complete, the remaining growth from each secondary plate was archived at −80°C in sterile STGG. Refer to Appendix 1 for a more detailed description of latex serotyping.

### Microarray Serotyping

For samples with culture-confirmed pneumococcal carriage, the original inoculated STGG was thawed and vortexed. Aliquots of 100μl were shipped in 1.8mL cryovials to BUGS Bioscience (BUGS Bioscience Ltd., London, United Kingdom) on dry ice. (Figure 1) The remaining steps for microarray serotyping (including sample processing, culturing, DNA extraction, molecular serotyping and analysis) were completed entirely by BUGS Bioscience. (16, 17) Final microarray results were retrieved by the study team from BUGS Bioscience’s web-based SentiNet platform and imported into STATA 13.1 (StataCorp, College Station, TX, USA) for analysis. Refer to Appendix 1 for a more detailed description of microarray serotyping.

### DNA Extraction and WGS

Archived secondary growth isolates were used to develop sequence libraries for serotyping-by-sequencing (via PneumoCat). To optimise total retrieved DNA, 30μl of thawed isolate-STGG was incubated overnight in 6mL THY (Todd Hewitt broth + yeast) enrichment culture. DNA was extracted from the overnight culture using the Qiagen^®^ QIAamp™ DNA Mini Kit, following manufacturer guidelines for bacterial DNA. Quality control (QC) measures, as required by the guidelines of the sequencing institution, included DNA quantification (Qubit™, Thermo Fisher Scientific, Massachusetts, USA) of all DNA samples and gel electrophoresis imaging on 0.7% agarose to assess DNA integrity. After attaining quantity and quality requirements, 100μL of extracted DNA were aliquoted into skirted 96-well microwell plates and stored at −80°C until shipped on dry ice to the Oxford Genomics Centre (University of Oxford, United Kingdom) for sequencing. Whole genome sequencing was performed at the Oxford Genomics Centre on a HiSeq4000 platform (Illumina™), with paired-end libraries and a read length of 150 pb.

### Serotyping-by-sequencing

WGS data was retrieved by the study team from a web-based FTP link. Serotype was inferred from the isolates’ genome sequences using the PneumoCaT software pipeline, an opensource bioinformatic tool. (18) PneumoCaT requires raw sequencing reads for each isolate which were trimmed and cleaned. Reads were trimmed of the illumina adapters and cleaned of low-quality ends using Trimmomatic (ver. 0.38; available at http://www.usadellab.org/cms/?page=trimmomatic). Minimum read length after trimming was 80 base pairs (bp), with the minimum average quality for a sliding window of 4 nucleotide being 15. A subset of 700,000 reads per end (1.4 million total) was used for any subsequent analysis. XML result files were parsed with ad hoc bash scripts, in order to extract and tabulate the serotyping result for each isolate. PneumoCaT was installed and used on a Linux machine at the MRC Cloud Infrastructure for Microbial Bioinformatics (CLIMB; https://www.climb.ac.uk/). Each serotype identification required an average 5-8 minutes. Refer to Appendix 1 for a more detailed description of serotyping-by-sequencing.

### Definitions

Concordance was calculated with all samples aggregated and according to the level of discrimination provided by the method. Concordance is reported using two criteria: i) a criterion based on whether both assays reported NVT or both reported VT (VT/NVT criterion) and ii) a criterion based on whether the final serotype reported by each assay is equivalent (serotype-specific criterion).

Concordance between latex and serotyping-by-sequencing (PneumoCaT): Other than a limited number of serogroups (6, 7, 9, 18, 19, 23) for which the latex kit provides serogroup differentiation, there is no further differential identification of NVT serogroups to serotype. NVT and non-typeable isolates were reported as NVT. Concordance at serotype level (serotype-specific criterion) was reported only if latex reported VT carriage. If latex reported NVT, any NVT reported by PneumoCaT was considered concordant. For example: 23F reported by both latex and PneumoCaT was considered concordant, as was NVT and 15B. However, 19F and 19A was considered discordant, as was NVT and 6B.

Concordance between latex and microarray: Concordance at serotype level (serotype-specific criterion) was reported only if latex reported VT carriage. If latex reported NVT, any NVT reported by microarray was considered concordant. Because microarray reports multiple serotype carriage, 23F reported by latex and 23F+34 reported by microarray was considered concordant, as was NVT and 18C+33D. However, 19F and 33D+19A was considered discordant, as was NVT and 3+7F. Note that for microarray, some closely related serotypes were reported as a group, with the individual serotype call in brackets (e.g., 6A/B [6B]). In this case, results were analysed to the level of the individual serotype call. For simplicity of analysis, if a method did not claim to detect a serotype (e.g. 23F) but the sample contained that serotype, this result was deemed discordant.

Concordance between microarray and serotyping-by-sequencing (PneumoCat): Microarray and PneumoCat both differentiate VT and NVT to serotype level, allowing concordance to be calculated on serotype concordance (serotype-specific criterion) for both VT and NVT *S. pneumoniae*.

### Statistical analysis

The formula for percent increase in VT prevalence was: ([VT prevalence using latex – VT prevalence using microarray] / VT prevalence using latex) * 100%. Confidence intervals are binomial exact. Statistical significance was inferred from two-sided p<0.05. Participant data collection was completed using Open Data Kit (ODK) Collect open source software. (v1.24.0). Statistical analyses were completed using Stata 13.1 (StataCorp, College Station, TX, USA).

### Ethics Considerations

The study protocol was approved by the College of Medicine Research and Ethics Committee, University of Malawi (P.02/15/1677) and the Liverpool School of Tropical Medicine Research Ethics Committee (14.056). Adult participants and parents/guardians of child participants provided written informed consent; children 8-years and older provided informed assent. This included consent for publication.

## Results

Pneumococcal carriage prevalence results from the larger surveillance project are reported elsewhere. (14) Comparing latex and PneumoCat, the adjusted concordance of correctly identifying pneumococcal carriage as VT or NVT was 90.7% (1216/1341; 95% CI: 89.0– 92.2). (Figure 3) Based on the serotype-specific criterion, concordance between latex and PneumoCaT was 87.5%; (1174/1341) (95% CI: 85.7–89.3). Comparing latex and microarray, the concordance based on correctly identifying pneumococcal carriage as VT or NVT was 97.3% 1311/1347 (95% CI: 96.3–98.1). Based on a serotype-specific criterion, the concordance was 95.2% (1282/1347; 95% CI: 93.9–96.3). Comparing microarray and PneumoCaT, concordance based on correctly identifying pneumococcal carriage as VT or NVT was 96.6% (1295/1341; 95% CI: 95.5–97.5). Based on a serotype-specific criterion, the concordance was 82.8% (1110/1341; 95% CI: 80.6–84.8).

**Figure 3.**
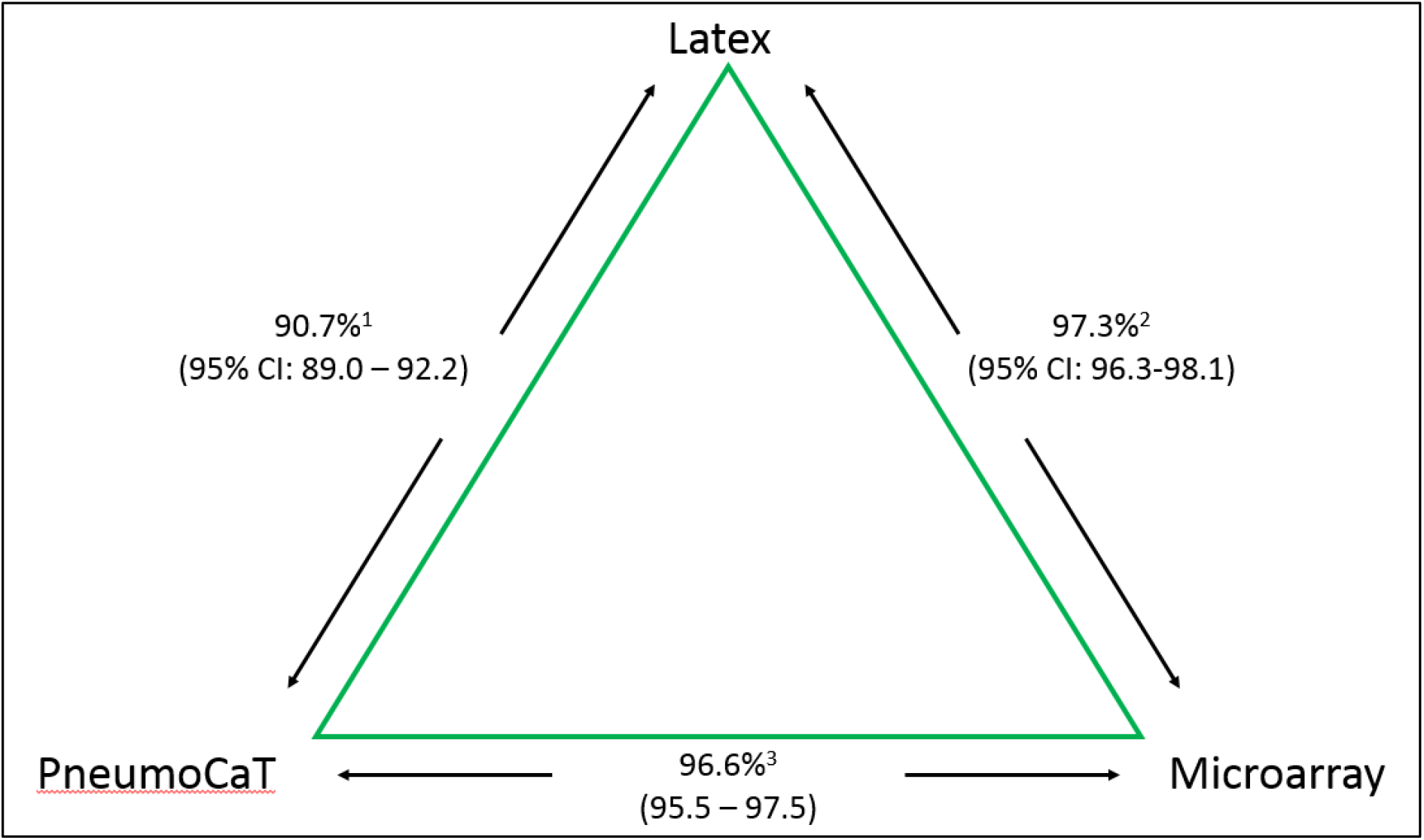
Concordance between assays. Concordance between two assays was defined as both assays identifying pneumococcal carriage as VT or both as NVT. Latex and PneumoCaT reported one result per sample, both using the same pure-growth culture. Microarray, using an aliquot of the original NPS-STGG, differentiated individual serotypes in multiple serotype carriage, when present. When comparing the three assays, concordance was based on serotype if latex reported VT carriage. If latex reported NVT, this was considered concordant to any NVT reported by PneumoCaT and microarray, as long as PneumoCaT and microarray reported the same NVT.

### Increased VT Detection Using Microarray

Using a larger study database of 1,949 samples from the same study, we evaluated latex and microarray data. Aggregating all ages (i.e. child and adult), there was an increase of 31.5% in VT prevalence by microarray compared to latex serotyping: 43.0% increase in VT carriage among children 3–6 years old, 21.7% among children 5–10 years old and 10.8% among HIV-infected adults on ART (Table 1). This was due to samples reporting NVT by latex but that also carried VT, as detected by microarray. These VT, undetected by latex, were carried in lower relative abundance (median 8%, range: 2% - 48%). The prevalence of multiple serotype carriage (range 2-6 serotypes) was 35.2% (686/1949). The prevalence among respective age groups was 44.4% (457/1029), 32.8% (169/515), and 14.8% (60/405). Among samples with multiple serotype carriage, latex identified the dominant serotype in 85.3% (585/686; 95%CI: 82.4–87.8) of samples. Despite the overall increase in detection of VT carriage, the proportion of individual VT serotypes detected is not different when comparing microarray to latex (Figure 4).

**Table 1.** Increased detection of VT carriage, latex vs microarray

|  | Latex<br>VT prevalence (n)<br>95% CI | Microarray<br>VT prevalence (n)<br>95% CI | % Increase<br>in VT prevalence |
| --- | --- | --- | --- |
| Children 3-6 years,<br>PCV-vaccinated (n=1360) | 20.0% (272)<br>17.9, 22.2 | 28.6% (389)<br>26.2, 31.1 | 43.0% |
| Children 5-10 years,<br>PCV-unvaccinated (n=904) | 21.1% (191)<br>18.5, 23.9 | 26.5% (240)<br>23.7, 29.6 | 21.7% |
| Adults, 18-40 years, HIV-infected,<br>PCV-unvaccinated (n=963) | 14.2% (137)<br>12.1, 16.6 | 16.6% (160)<br>14.3, 19.1 | 10.8% |
| Total (n=3227) | 18.6 (600)<br>17.3, 20.0 | 24.4 (789)<br>23.0, 26.0 | 31.5% |

PCV=pneumococcal conjugate vaccine, VT=vaccine serotype, CI=confidence interval.

**Figure 4.**
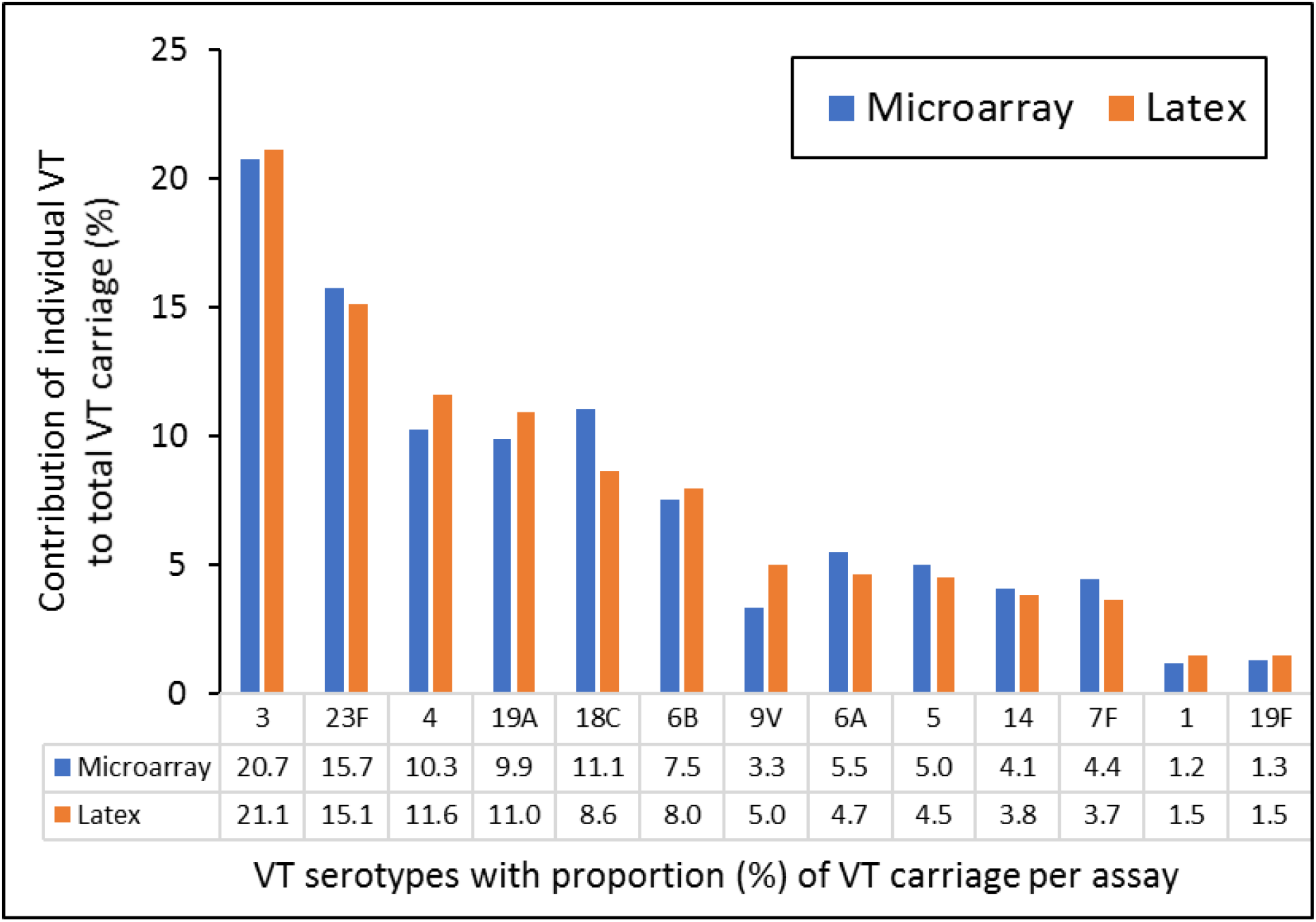
Proportion of individual VT serotypes contributing to total VT carriage. The proportion of individual VT serotypes detected is not significantly different when comparing microarray to latex. VT=vaccine serotype.

### Key Parameters of selected serotyping methods

Table 2 presents key parameters to further consider when deciding which assay is appropriate for a particular setting. Estimated costs and feasibility of implementation and maintenance are specific to the setting in Malawi at the Malawi-Liverpool-Wellcome Trust Clinical Research Programme in Blantyre. Extrapolation would need further validation outside the scope of this evaluation. Though more limited in its reporting only a single serotype, latex is highly accurate while being less costly and requiring less expertise and resources for field-implementation and analysis. While microarray is the costliest option, it provides greater accuracy of total pneumococcal carriage, including multiple serotype carriage and relative abundance of individual serotypes in carriage. Whole genome sequencing is a strong alternative to latex and would be nearly cost-free if the sequence libraries were already available. In addition, WGS provides opportunity for further analyses, including population structure and antibiotic resistance.

**Table 2.**
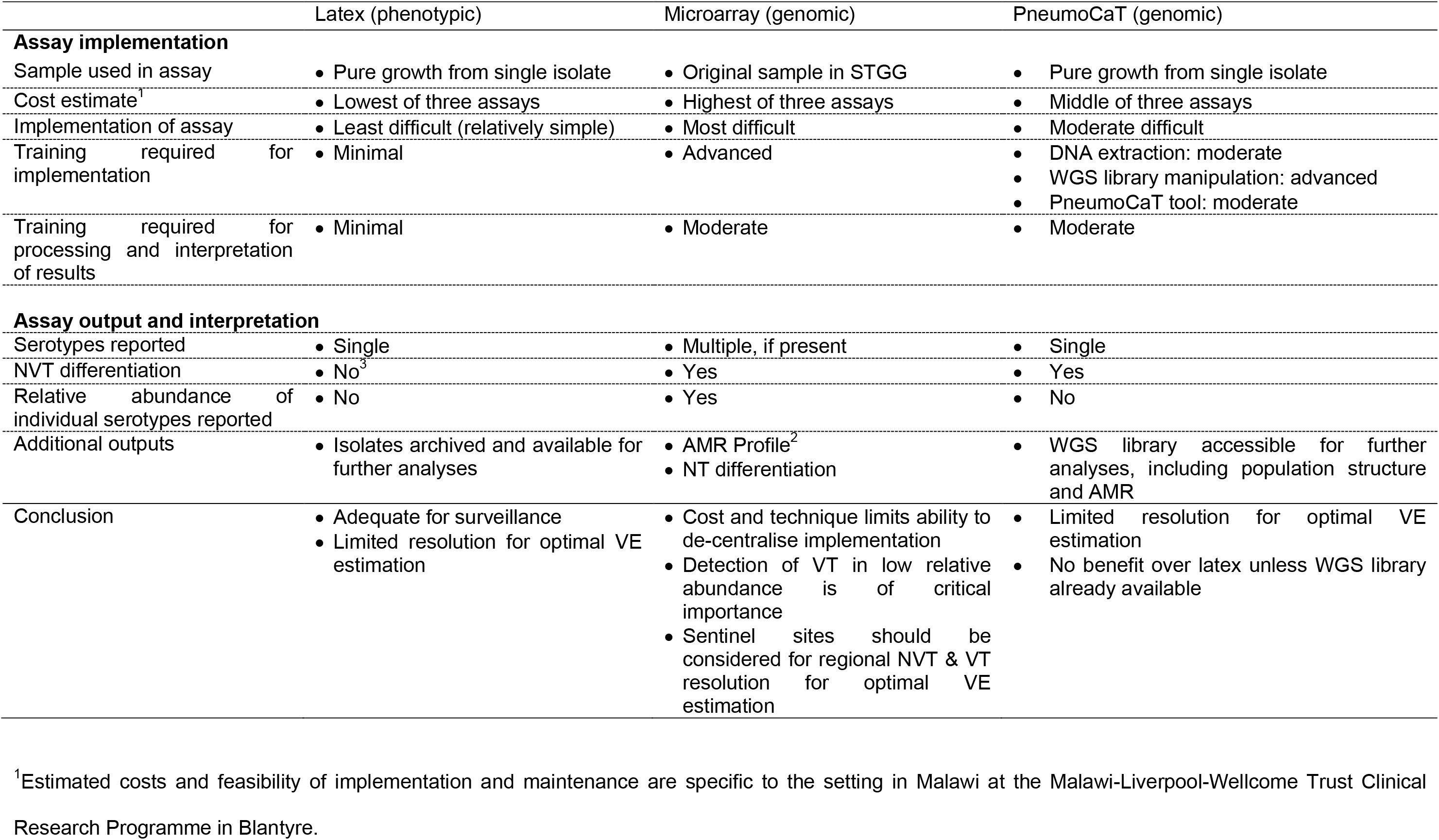
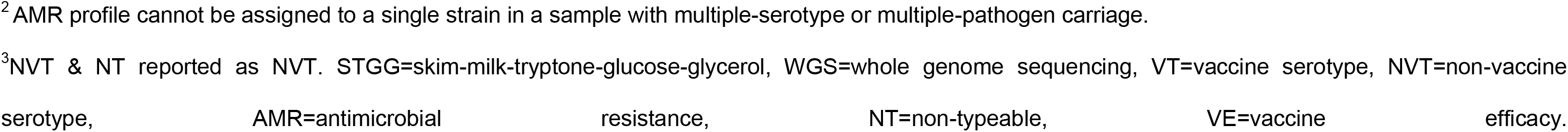
Key comparative parameters of serotyping methods.

|  | Latex (phenotypic) | Microarray (genomic) | PneumoCaT (genomic) |
| --- | --- | --- | --- |
| **Assay implementation** |  |  |  |
| Sample used in assay | • Pure growth from single isolate | • Original sample in STGG | • Pure growth from single isolate |
| Cost estimate<sup>1</sup> | • Lowest of three assays | • Highest of three assays | • Middle of three assays |
| Implementation of assay | • Least difficult (relatively simple) | • Most difficult | • Moderate difficult |
| Training required for implementation | • Minimal | • Advanced | • DNA extraction: moderate<br>• WGS library manipulation: advanced<br>• PneumoCaT tool: moderate |
| Training required for processing and interpretation of results | • Minimal | • Moderate | • Moderate |
| **Assay output and interpretation** |  |  |  |
| Serotypes reported | • Single | • Multiple, if present | • Single |
| NVT differentiation | • No<sup>3</sup> | • Yes | • Yes |
| Relative abundance of individual serotypes reported | • No | • Yes | • No |
| Additional outputs | • Isolates archived and available for further analyses | • AMR Profile<sup>2</sup><br>• NT differentiation | • WGS library accessible for further analyses, including population structure and AMR |
| Conclusion | • Adequate for surveillance<br>• Limited resolution for optimal VE estimation | • Cost and technique limits ability to de-centralise implementation<br>• Detection of VT in low relative abundance is of critical importance<br>• Sentinel sites should be considered for regional NVT & VT resolution for optimal VE estimation | • Limited resolution for optimal VE estimation<br>• No benefit over latex unless WGS library already available |

<sup>1</sup>Estimated costs and feasibility of implementation and maintenance are specific to the setting in Malawi at the Malawi-Liverpool-Wellcome Trust Clinical Research Programme in Blantyre.
<sup>2</sup> AMR profile cannot be assigned to a single strain in a sample with multiple-serotype or multiple-pathogen carriage.
<sup>3</sup>NVT & NT reported as NVT. STGG=skim-milk-tryptone-glucose-glycerol, WGS=whole genome sequencing, VT=vaccine serotype, NVT=non-vaccine
serotype, AMR=antimicrobial resistance, NT=non-typeable, VE=vaccine efficacy.

## Discussion

We report high concordance between three serotyping techniques applicable to routine pneumococcal surveillance. Importantly, we have extended the analysis to include relevant parameters beyond accuracy including cost, time to result, and measures of input required for assay implementation and maintenance. These are parameters that researchers and policy makers should consider when deciding which assay to implement. All three assays appear accurate and concordant in identifying the dominant serotype.

While latex agglutination is accurate and requires the least expertise and resources for field-implementation and analysis and provides rapid results, standard latex approaches is not optimal for optimal surveillance of vaccine impact, including the detection of multiple serotype carriage and VT in low relative abundance. (19) There have been attempts to implement latex for detection of multiple serotype carriage. Gratten et al. serotyped up to six colonies from nasal-swab culture plates and found multiple-serotype carriage in 29.5% of Papua New Guinean children. (20) The authors went on to serotype at least 50 colonies from 10 selected nasal-swab cultures and concluded that the minor carried serotype accounted for 4 to 27% of the total pneumococcal population. A review of published data on multiple carriage concluded that, to detect a minor carried serotype it would be necessary to serotype at least five colonies to have a 95% chance of detecting the serotype if it accounted for 50% of the total pneumococcal population, and one would need to examine 299 colonies if the serotype was present at a relative abundance of 1%. As part of the PneuCarriage project, to thoroughly characterise samples, up to 120 colonies from each sample were selected to achieve >99% power to detect a minor serotype of 5% abundance. (13) This approach would not be cost- or time-effective. Though dependent on technical capacity to develop in-house reagents, researchers in The Gambia developed a latex agglutination technique in which colonies from the primary culture plate are suspended in saline and serotyped by latex agglutination. (21) While not differentiating NVT serotypes, they did show that up to 10.4% of pneumococcal acquisitions were found to be of multiple serotypes in a longitudinal infant cohort study. While latex is limited in its output, the process can be leveraged for additional endpoints including, for example, measuring carriage density through counting of colony-forming units (CFU) on agar culture plates.

With opensource bioinformatic tools such as PneumoCaT, serotyping-by-sequencing can be less costly than microarray, even accounting for costs of DNA extraction and WGS, while still being able to differentiate non-typeable and nearly every known VT and NVT. Though we would not recommend initiating DNA extraction and WGS for the use of PneumoCaT alone, sequence libraries can be further leveraged for extensive informative bioinformatic analyses, useful in population biology, antimicrobial resistance investigations and vaccine monitoring. Moreover, using PneumoCaT for serotyping would be essentially cost-free if the sequence libraries were already available, apart from the limited bioinformatic skills needed. While microarray is more costly, it differentiates NVT and multiple serotype carriage with relative abundance, as well as non-*S. pneumoniae* contaminants (i.e. *S. mitis, S. salivarius*, *Staphylococcus aureus*) with a degree of precision. This technique stands out for its sensitivity, being able to detect serotypes in low relative abundance, which is of critical importance for understanding the transmission patterns of *S. pneumoniae*.

There are a number of limitations to mention, including the number of serotyping methods which were not evaluated, including PCR and the SeroBA pipeline. SeroBA is a relatively new serotyping-by-sequencing software. With similar accuracy to PneumoCaT, SeroBA does have operational advantages. (22) SeroBA can correctly call a serotype with a read coverage as low as 10X (20X is required for PneumoCaT). Using a k-mer based approach, rather than the raw sequence alignment, SeroBA requires much lower computational power and time. On the other hand, the PneumoCat source code can be easily adapted to the operator needs, and both software are likely to run on a standard server configuration. Alternative culture-independent methods, such as PCR, could be important for confirming carriage when re-culturing of original NP swab samples is not feasible. Though PCR has been successfully applied on DNA extracted directly from NPS-STGG, evidence suggests that the best way to apply PCR serotyping is after culture enrichment, returning a higher sensitivity and ability to identify multiple serotype carriage. (9) PCR limitations include the need for region-specific reaction protocols, implementing a high number of primer pairs to identify the same range of serotypes identified by microarray or WGS, and the increased risk of detecting non-viable pneumococci. As there is no evidence of a viable but non-culturable (VBNC) state in *S. pneumoniae*, (23) identifying non-viable pneumococci could be disadvantageous for field-based research. While a formal economic analysis of the methods would be justified, we were unable to extrapolate the individual costing components between sites. Such components would include local salaries and additional labour costs, procurement and shipping of equipment and consumables, equipment maintenance, local health and safety requirements, and institutional costs. For this reason, comparative costing is grossly categorized. Though we did not include invasive isolates (from blood or cerebral spine fluid, for example), it is important to identify serotypes associated with IPD, including in post-PCV impact studies. For invasive isolates, with a single-serotype sample, microarray would have limited advantage. Application of serotyping-by-sequencing would then be the most informative option, including insight into population structure, antimicrobial resistance patterns and serotype replacement disease.

## CONCLUSION

Selection of the appropriate assay should be based on the intended analysis and endpoint. While accuracy and concordance is high between the three assays, parameters of field-implementation and cost vary significantly. In a setting of limited resources, as is true throughout much of sub-Saharan Africa, latex is the best overall option for decentralised surveillance of vaccine impact. However, WGS, which adds population structure, and microarray, which adds multiple-serotype carriage, should be considered at regional reference laboratories while investigating the importance of VT in low relative abundance in transmission and disease.

## Appendixes

An appendix is available online. Consisting of data provided by the authors to benefit the reader, the posted materials are not copyedited and are the sole responsibility of the authors, so questions or comments should be addressed to the corresponding author.

## Data availability

The data supporting the findings of this study has been deposited in the Figshare repository. (24)

## Acknowledgements

The authors thank the individuals who participated in this study and the local authorities for their support. We are grateful to the hospitality of the QECH ART Clinic, led by Ken Malisita. Our thanks also extend to the MLW laboratory management team, led by Brigitte Denis and George Selemani.

## Notes

### Disclaimer

The funders had no role in study design, collection, analysis, data interpretation, writing of the report or in the decision to submit the paper for publication. The corresponding author had full access to the study data and, together with the senior authors, had final responsibility for the decision to submit for publication. TDS, AG, RSH, and NF are supported by the National Institute for Health Research (NIHR) Global Health Research Unit on Mucosal Pathogens using UK aid from the UK Government. The views expressed in this publication are those of the author(s) and not necessarily those of the NIHR or the Department of Health and Social Care.

